# *Drosophila scribble* mutant tumors undergo a transition from a growth arrest state to a proliferative state over time

**DOI:** 10.1101/450486

**Authors:** Tiantian Ji, Lina Zhang, Shengshuo Huang, Mingxi Deng, Ying Wang, Tri Thanh Pham, Clemens Cabernard, Jiguang Wang, Yan Yan

## Abstract

The *Drosophila* neoplastic tumor suppressor gene (nTSG) mutant tumors have successfully modeled many aspects of human tumor progression. However, the fly nTSG mutant tumors progress rapidly over days. This is in contrast with most human tumors which develop slowly, harbor heterogeneous cell populations for selection and undergo an evolution-like process. Whether the fast-growing fly nTSG mutant tumors have capacity for evolution remains unclear. Through quantitative analysis of the *scrib* mutant tumor growth, we found that the *scrib* mutant tumors evolve to display different growth rates and cell cycle profiles over time. Multiple growth-regulatory signaling pathways show quantitative differences in early versus late *scrib* mutant tumors. These data suggest that the *scrib* mutant tumors undergo a transition from a growth arrest state to a proliferative state. Through longitudinal single cell RNA (scRNA) data analysis we found that the *scrib* mutant tumors harbor heterogeneous cell populations likely of distinct proliferative states, which are available for potential selection. This study raises the possibility of studying tumor evolution in a genetically accessible and fast-growing invertebrate tumor model.

## Introduction

Proteins essential for maintaining epithelial structures, such as cell polarity complexes, are involved in growth control (Bilder, 2004; Boggiano and Fehon, 2012; Sun and Irvine, 2016). For example, the basolateral Scribble complex, composed of Scribble (Scrib), Discs large (Dlg) and Lethal giant larvae (L(2)gl), were discovered as a group of “neoplastic tumor suppressor genes” (nTSGs) in *Drosophila* (Bilder et al., 2000; Bilder and Perrimon, 2000; Gateff, 1978; Woods and Bryant, 1991). *Drosophila* larvae homozygous mutant for any of the nTSGs grow into giant larvae with tumorous imaginal discs and optic lobes. These mutant tumors fail to differentiate and grow into masses that survive serial transplantations, induce cachexia and eventually kill the hosts (Figueroa-Clarevega and Bilder, 2015; Gateff, 1978). Studies of *Drosophila* nTSGs over decades have provided valuable insights into the mechanisms of growth control and tumorigenesis (Bilder, 2004; Gonzalez, 2013; Pastor-Pareja and Xu, 2013; Richardson and Portela, 2018; Sonoshita and Cagan, 2017). For example, analyses of the nTSG mutant clonal growth have revealed cell competition-mediated tumor suppression mechanisms, the cooperative actions of multiple conserved signaling pathways during tumor development, and tumor microenvironment influences (Brumby and Richardson, 2003; Chen et al., 2012; Cordero et al., 2010; Igaki et al., 2006; Katheder et al., 2017; Pagliarini and Xu, 2003; Vaughen and Igaki, 2016; Yamamoto et al., 2017).

Interestingly, while the fly nTSG mutant tumors have successfully modeled many aspects of human epithelial cancers, it was noted that for the fly nTSG tumors that progress rapidly over days, a single gene mutation is sufficient to cause tumorigenesis. This is in contrary to human tumors which typically develop slowly over months and years and supported a multiple-hit model (Bilder, 2004; Hanahan and Weinberg, 2000; Nordling, 1953). Human tumors have been shown to display a variable degree of genetic and epigenetic intratumor heterogeneity (ITH) that provides a foundation for selection and tumor evolution (McGranahan and Swanton, 2017). Whether the fast-growing fly nTSG mutant tumors have capacity for evolution has remained unclear.

Through quantitative analysis of the *scrib* mutant tumor growth, we found that over time the *scrib* mutant tumors display different growth rates and cell cycle profiles. Moreover, multiple signaling pathways display quantitative differences in early versus late *scrib* mutant tumors. We demonstrated that high JNK signaling activity is a primary cause of growth arrest in early *scrib* mutant tumors. These data suggest that the *scrib* mutant cells undergo a transition from a growth arrest state to a proliferative state during tumor progression. Longitudinal scRNA data analysis further reveals heterogeneity in the *scrib* mutant tumors that potentially provides opportunities for selection and drives the transition from a growth arrest state to a proliferative state as a population.

## Results

### The *scrib* mutant tumors display different growth rates over time

To explore potential evolving traits during the *scrib* mutant tumor progression, we first monitored the growth of wing imaginal discs derived from 3-hour egg collection of a *scrib^1^*/TM6B stock (Bilder and Perrimon, 2000) (Figure 1A-B). On average, at 4-day and 5-day after egg laying (AEL), the growth rate of the *scrib* mutant tumors is around 25%-30% of that of control imaginal discs raised at identical conditions (Figure 1B). By 7-day AEL the growth rate of the *scrib* mutant tumors is comparable with that of the 5-day AEL control group (Figure 1B). Note that the *scrib* mutant tumors would continue growth to sizes consistent with previous reports (Bilder et al., 2000; Gateff, 1978). Using phospho-Histone H3 (PH3) as a marker for mitotic cells, we detected the 4-day AEL *scrib* mutant tumors harbor much less PH3+ cells per unit volume than the control group (Figure 1C-D and 1G). By 5-day AEL the *scrib* mutant tumors contain comparable number of PH3+ cells per unit volume with the control larvae as the wild type control group approached the end of growth period (Figure 1E-F and 1G).

**Figure 1.**
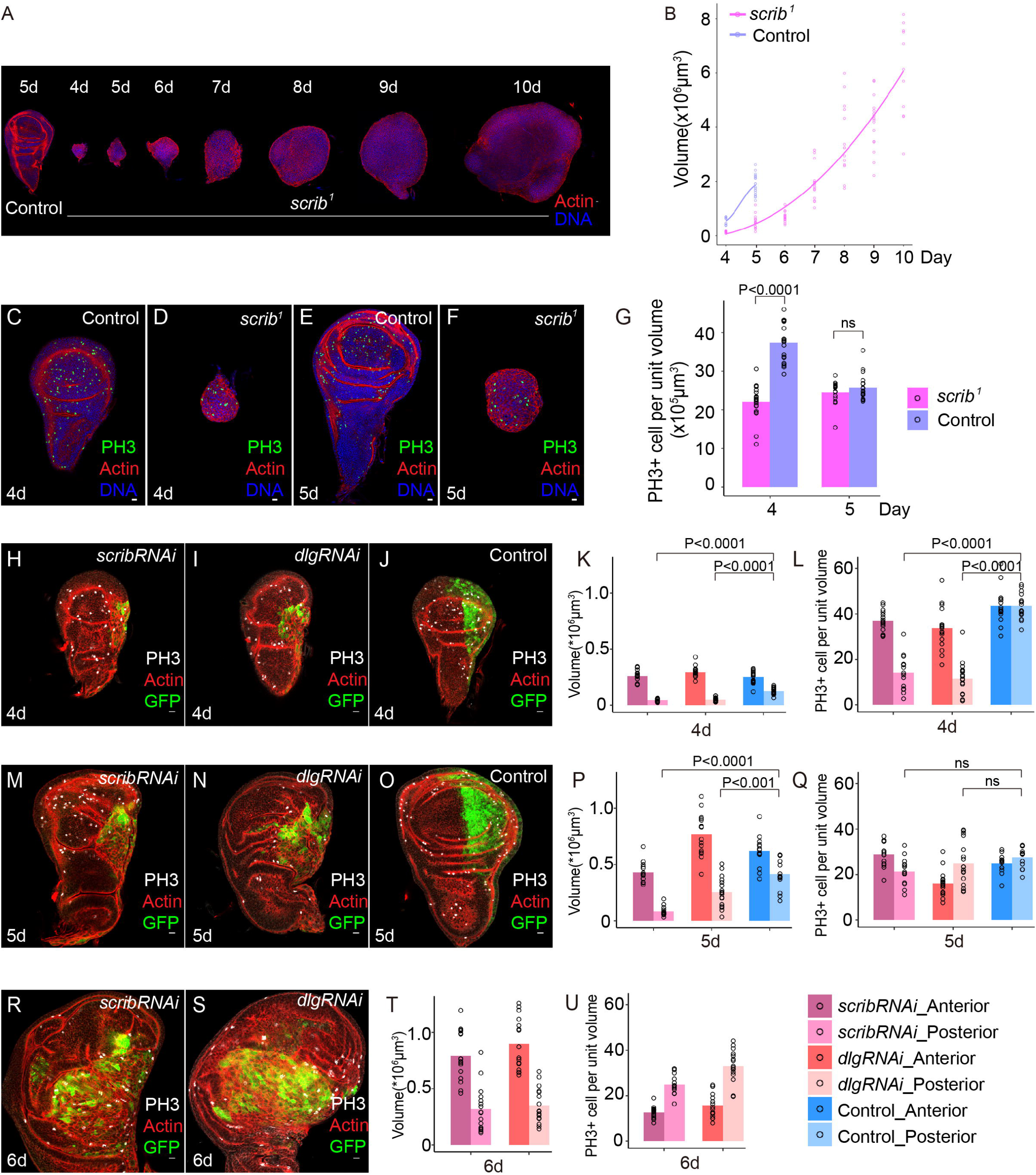
The Scrib-depleted cells display different growth rates over time. (A) Examples of a control 5-day AEL imaginal disc and *scrib^1^* mutant wing imaginal discs from 4-day AEL to 10-day AEL stained for actin (red) and DNA (blue). Control genotype: *FRT82B*. Scale bar: 10μm. (B) Quantification of volumes for control and *scrib^1^* mutant wing imaginal discs over time. Control genotype: *FRT82B* raised at identical conditions. Control, **4d** n = 10, 5±1×10^5^μm^3^, **5d** n =19, 1.9±0.4×10^6^μm^3^; *scrib^1^* mutant, **4d** n = 16, 1.4±0.3×10^5^μm^3^, **5d** n = 18, 5±2×10^5^μm^3^, **6d** n = 19, 7±2×10^5^μm^3^, **7d** n = 17, 2.0±0.7×10^6^μm^3^, **8d** n = 14, 3±1×10^6^ μm^3^, **9d** n = 17, 4±1×10^6^ μm^3^, **10d** n = 11, 6±2×10^6^ μm^3^. Note that larvae from the control group become pupae at 5-day AEL. (C-F) 4-day AEL (C-D) and 5-day AEL (E-F) control (C and E) and *scrib^1^* mutant (D and F) wing imaginal discs stained for PH3(green), actin (red) and DNA (blue). Control genotype: *FRT82B*. Scale bar: 10μm. (G) Quantification of PH3+ cell number per unit volume (10^5^μm^3^) in 4-day AEL and 5-day AEL control and *scrib^1^* mutant wing imaginal discs. Statistical analysis was performed by unpaired t-test. Control, *FRT82B*, **4d** n = 10, 37±5, **5d** n =19, 26±3; *scrib^1^* mutant, **4d** n = 16, 22±4, **5d** n = 18, 25±3. (H-U) 4-day (H-J), 5-day (M-O) and 6-day (R-S) AEL *scribRNAi* (H, M, R), *dlgRNAi* (I, N, S) and control (J, O) imaginal discs stained for PH3 (gray), actin(red) and GFP (green). Quantification of volumes (K, P, T) and PH3+ cell number per unit volume (10^5^μm^3^) (L, Q, U) for 4-day (K, L), 5-day (P, Q) and 6-day (T, U) AEL *scribRNAi*, *dlgRNAi* and control imaginal discs. Genotype for the *scribRNAi* group: *engrailed-Gal4 UAS-GFP/+; UAS-scribRNAi/+.* Genotype for the *dlgRNAi* group: *engrailed-Gal4 UAS-GFP/+; UAS-dlgRNAi/+.* Genotype for the control group: *engrailed-Gal4 UAS-GFP/+; P{y[+t7.7]=CaryP}attP2 /+* (the 3rd chromosome TRiP line background strain). The *scribRNAi* group, **4d** n = 15, anterior, 2.6±0.5×10 μm, PH+ cell number 37±5, posterior, 4±1×10^4^μm^3^, PH+ cell number 14±8, **5d** n = 14, anterior, 4.3±0.9×10^5^μm^3^, PH+ cell number 29±5, posterior, 8±4×10^4^μm^3^, PH+ cell number 21±6, **6d** n = 15, anterior, 8±2×10^5^μm^3^, PH+ cell number 13±3, posterior, 3±2×10^5^μm^3^, PH+ cell number 25±4; The *dlgRNAi* group, **4d** n = 16, anterior, 3.0±0.5×10 μm, PH+ cell number 34±9, posterior, 5±2×10^4^μm^3^, PH+ cell number 11±8, **5d** n = 17, anterior, 8±2×10^5^μm^3^, PH+ cell number 16±5, posterior, 3±1×10^5^μm^3^, PH+ cell number 25±9, **6d** n = 16, anterior, 9±2×10^5^μm^3^, PH+ cell number 16±5, posterior, 4±2×10^5^μm^3^, PH+ cell number 33±7; The control group, **4d** n = 16, anterior, 2.5±0.6×10 μm, PH+ cell number 44±8, posterior, 1.3±0.3×10^5^μm^3^, PH+ cell number 43±6, **5d** n = 13, anterior, 6±1×10^5^μm^3^, PH+ cell number 25±4, posterior, 4±1 x10^5^μm^3^, PH+ cell number 28±4. Scale bar: 10μm. Statistical analysis was performed by unpaired t-test. Note that larvae from the control group become pupae at 5-day AEL.

To test whether the initial slow-growth phenotype we observed in the early *scrib* mutant tumors is specific to the *scrib^1^* allele, we generated tumors depleted of Scrib or Dlg at the posterior region of wing discs through *engrailed*-Gal4 mediated RNAi. Although the mosaic clones depleted of Scrib or Dlg are eliminated through cell competition (Brumby and Richardson, 2003; Igaki et al., 2006; Vaughen and Igaki, 2016; Yamamoto et al., 2017), the cell competition process does not cross segmentation boundary (Johnston, 2009; Morata and Ripoll, 1975; Simpson, 1979; Simpson and Morata, 1981). Therefore, we can analyze the growth of the posterior *scrib RNAi* and *dlg RNAi* tumors independent of the influence of cell competition. At 4-day and 5-day AEL, the volume of the *scrib* RNAi and *dlg* RNAi tumors is much smaller than the size of the posterior region in control wing discs (Figure 1H-Q). Meanwhile, the anterior regions of the *scrib* RNAi, *dlg* RNAi and control imaginal discs have comparable average volumes (Figure 1H-Q), indicating that the slow-growth phenotype observed in early *scrib* tumors is likely to be independent of specific alleles used and an overall larval developmental delay. Interestingly, from 4-day to 6-day AEL we could detect an increase of PH3+ cell number in the posterior *scrib* RNAi and *dlg* RNAi tumors (Figure 1H-U), indicative of changes in growth rates during tumor progression. Note that larvae harboring *scrib* RNAi or *dlg* RNAi imaginal discs turned into pupae by 7-day AEL, preventing further measurement of growth rates.

### The *scrib* mutant tumors show cell cycle defects that resolve over time

The small volume of early *scrib* mutant tumors can be caused through increased apoptosis, defects in cellular growth or defective cell proliferation. We found that prevention of apoptosis by overexpressing p35 could not rescue the growth arrest of the posterior *scrib* RNAi cells (Figure S1A-H). Moreover, while we were able to detect apoptotic cells in the posterior *scrib* RNAi region, a similar number of apoptotic cells can also be detected in the anterior control region (Figure S1I). Therefore, the small volume of early *scrib* mutant tumors is unlikely to be caused by increased apoptosis. The cell volume of individual *scrib* mutant cells is larger than that of the control wing disc cells (Figure S2A-B), consistent with a loss of epithelial packing and an elevation of mTOR signaling activity in the *scrib* mutant cells (Figure S2C-F). Therefore, the growth arrest of early *scrib* mutant tumors is also unlikely to be caused by defects in cellular growth.

Next, we examined additional proliferation markers in the *scrib* mutant tumors. Using 30-min EdU incorporation as an indicator for S-phase cells, we noticed a significant decrease in EdU incorporation in the 4-day AEL posterior *scrib* RNAi cells (Figure 2A, 2B and 2E). However, in 5-day AEL wing discs, we noticed that the posterior *scrib* RNAi cells show similar EdU incorporation rate as the control group (Figure 2C, 2D and 2E). Flow cytometry analysis showed a significant decrease of G1/S population in the posterior *scrib* RNAi cells in comparison with the anterior control cells from 4-day AEL wing discs (Figure 2F). By 5-day and 6-day AEL, the G1/S population in the posterior *scrib* RNAi cells showed progressive recovery to a level comparable with that of the anterior control cells (Figure 2G and 2H). We also observed a similar pattern of G1/S population decrease from 4-day AEL *scrib* mutant tumors and recovery from 5-day *scrib* mutant tumors in comparison with the control group (Figure 2I and 2J). Notably, flow cytometry analysis detected a population of cells with higher DNA content than normal cells in the 4-day AEL Scrib-depleted cells (Figure 2F and 2I). Using the FUCCI system (Zielke et al., 2014), we detected a population of G2/M cells (GFP+RFP+) with enlarged cell nuclei in 4-day AEL Scrib-depleted cells (Supplemental Movie 1). Taken together, the above data suggested that a population of 4-day AEL *scrib* mutant cells is arrested during G2/M transition and a subset of these cells might re-initiate the DNA replication process before entering mitosis.

**Figure 2.**
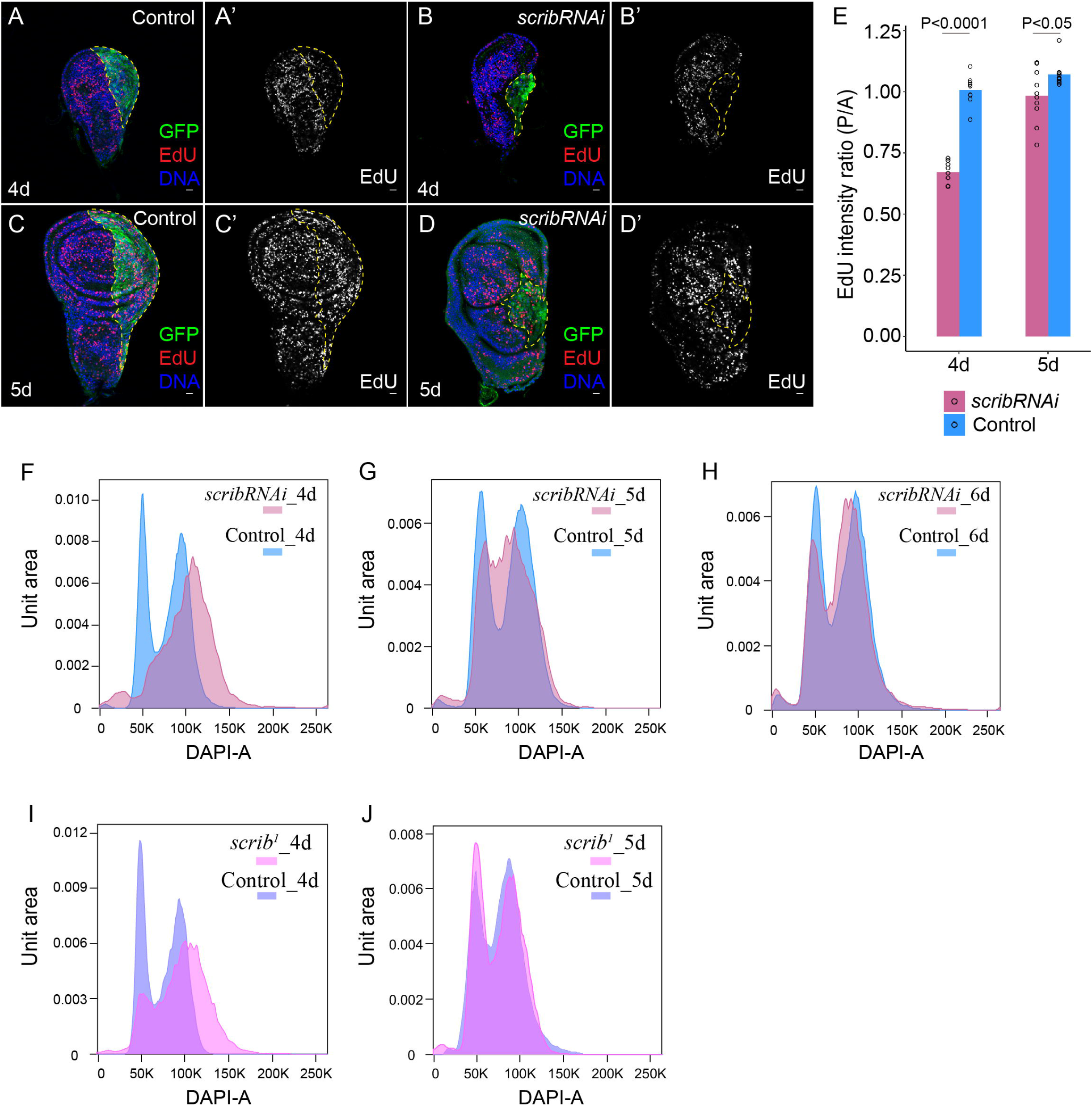
The Scrib-depleted cells display different cell cycle profiles over time. (A-E) 4-day AEL (A-B) and 5-day AEL (C-D) control (A and C) and *scribRNAi* (B and D) wing imaginal discs stained for EdU (red in A, B, C, D and gray in A’, B’, C’, D’), GFP (green) and DNA (blue). (E) Quantification of posterior/anterior EdU intensity ratio for 4-day and 5-day AEL control and *scribRNAi wing* imaginal discs. Genotype for the *scribRNAi* group: *engrailed-Gal4 UAS-GFP/+; UAS-scribRNAi/+.* Genotype for the control group: *engrailed-Gal4 UAS-GFP/+; P{y[+t7.7]=CaryP}attP2 /+.* The *scribRNAi* group, 4d n = 8, 5d n = 10; The control group, 4d n = 9, 5d n = 10. Statistical analysis was performed by unpaired t-test. Scale bar: 10μm. (F-H) FACS analysis of DNA contents of control and *scribRNAi* cells from 4-day (F), 5-day (G) and 6-day(H) AEL wing imaginal discs. Genotype for FACS analysis: *engrailed-Gal4 UAS-GFP/+; UAS-scribRNAi/+.* The anterior GFP- cells serve as control and the posterior GFP+ cells are the *scribRNAi* cells. Because the number of GFP- cells is much larger than GFP+ cells, the histogram overlay is normalized using the unit distribution mode in FlowJo. At least five thousand cells were recorded for each cell group. Each experiment is replicated for at least three times. (I-J) FACS analysis of DNA contents of control and *scrib^1^* mutant cells from 4-day (I) and 5-day (J) AEL wing imaginal discs. Genotype for the control group: *FRT82B* raised at identical conditions. At least five thousand cells were recorded for each cell group. Each experiment is replicated for at least three times.

*Drosophila* larval brain neuroblasts are an excellent model for analyzing mitosis due to its accessibility for live imaging (Cabernard and Doe, 2013). We observed that the *scrib* mutant neuroblasts displayed a significant prolonged entry into mitosis (Figure S3), consistent with the cell cycle defects we observed in 4-day AEL *scrib* mutant wing disc cells.

We conclude that the growth arrest of early *scrib* mutant tumors is most likely caused by defects in cell cycle progression, yet the cell cycle defects observed in the early *scrib* mutant tumors can resolve over time.

### The *scrib* mutant tumors display quantitative differences in multiple signaling pathway activities over time

To investigate the reason why early and late *scrib* mutant tumors display different growth rates and cell cycle profiles, we examined transcriptomes of *scrib* mutant tumors collected at different time points. Principal component analysis (PCA) showed the biological repeats of the *scrib* mutant tumors from the same stage are well-clustered and the *scrib* mutant tumors form a transition trajectory along time (Figure 3A). Hierarchical clustering further showed that the early and late *scrib* mutant tumors show distinctive gene expression pattern (Figure 3B).

**Figure 3.**
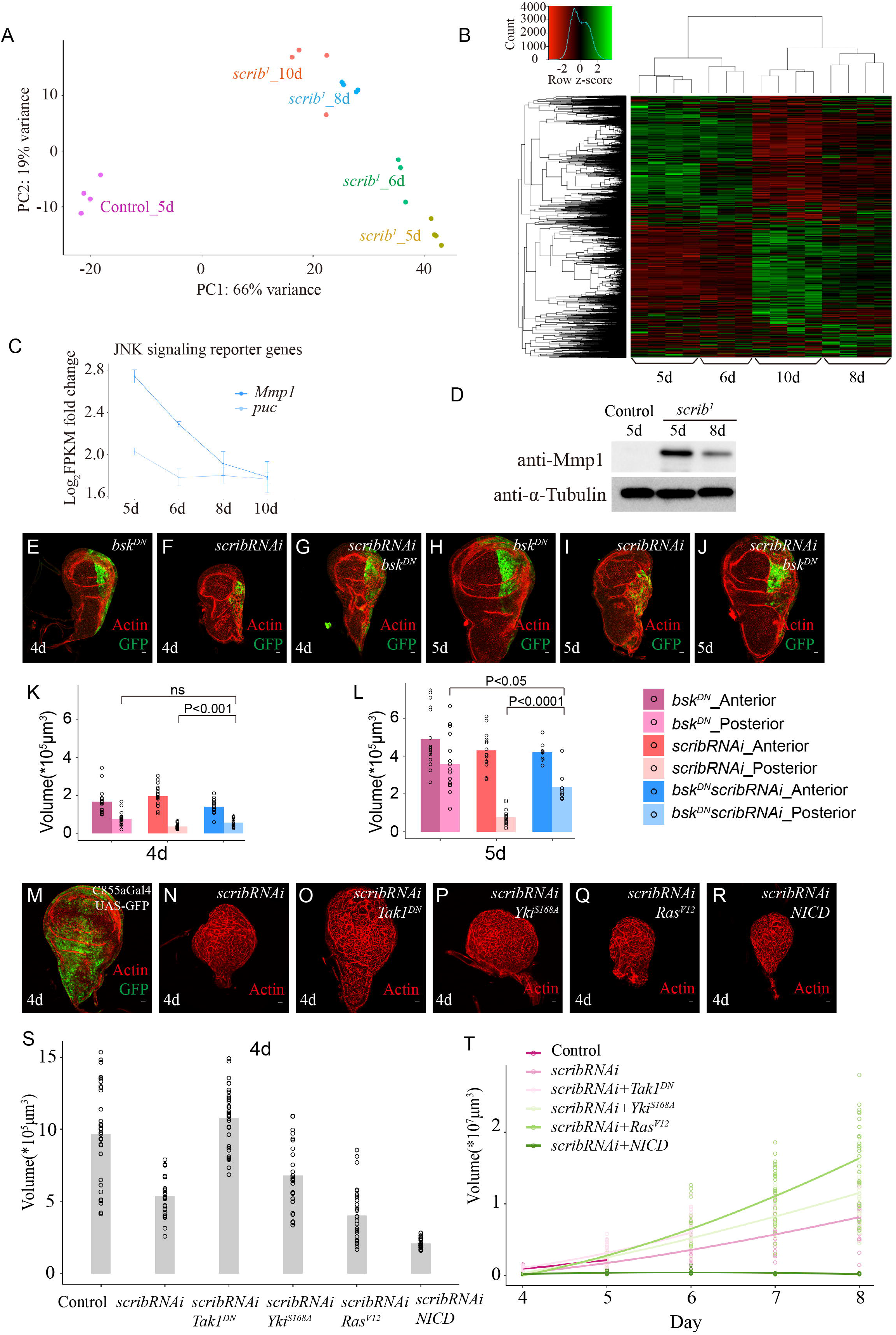
High JNK signaling activity causes growth arrest in early *scrib* mutant tumors. (A) Principle component analysis (PCA) of transcriptomes from 5-day AEL control imaginal discs and 5-day, 6-day, 8-day and 10-day AEL *scrib* mutant wing imaginal discs. Control genotype: *FRT82B*. Four biological replicates are plotted for each time point except for 6-day AEL groups which only three biological replicates are recovered. (B) Hierarchical clustering of transcriptomes from 5-day, 6-day, 8-day and 10-day AEL *scrib* mutant wing imaginal discs. (C) Plot of JNK signaling reporter genes in staged *scrib* mutant wing discs normalized by those in 5-day AEL control imaginal discs using log_2_(FPKM fold change) values. Control genotype: *FRT82B*. (D) Western blot analysis of Mmp-1 protein level in 5-day AEL control imaginal discs, 5-day and 8-day AEL *scrib* mutant wing imaginal discs. Control genotype: *FRT82B*. (E-J) 4-day (E-G) and 5-day (H-J) AEL *bsk^DN^* (E, H), *scribRNAi* (F, I) and *scribRNAi bsk^DN^* (G, J) imaginal discs stained for actin(red) and GFP (green). Genotype for (E) and (H): *engrailed-Gal4 UAS-GFP/+; UAS- bsk^DN^/+.* Genotype for (F) and (I): *engrailed-Gal4 UAS-GFP/UAS-scribRNAi.* Genotype of (G) and (J): *engrailed-Gal4 UAS-GFP/UAS-scribRNAi; UAS- bsk^DN^/+*. Scale bar: 10μm. (K-L) Quantification of volumes for 4-day (K) and 5-day (L) AEL *bsk, scribRNAi* and *scribRNAi bsk^DN^* imaginal discs. *bsk^DN^*, **4d** n = 17, anterior, 1.7±0.7×10^5^μm^3^, posterior, 8±4×10^4^μm^3^, **5d** n = 17, anterior, 5±2×10^5^μm^3^, posterior, 4±1×10^5^μm^3^. *scribRNAi*, **4d** n = 20, anterior, 2.0±0.5×10^5^μm^3^, posterior, 4±1×10^4^μm^3^, **5d** n = 15, anterior, 4±1×10^5^μm^3^, posterior, 8±4×10 μm. *scribRNAi bsk*, **4d** n = 14, anterior, 1.4±0.4×10 μm^3^, posterior, 6±2×10^4^μm^3^, **5d** n = 8, anterior, 4.2±0.5×10^5^μm^3^, posterior, 2.4±0.8×10^5^μm^3^. Statistical analysis was performed by unpaired t-test. (M-R) 4-day AEL control (M), *scribRNAi* (N)*, scribRNAi Tak1^DN^*(O*), scribRNAi Yki^S168A^* (P*), scribRNAi Ras^V12^(Q),* and *scribRNAi NICD* (R) imaginal discs stained for actin (red). Genotypes are as follows: (M) *C885a-Gal4/+; UAS-GFP/+;* (N) *C885a-Gal4/+; UAS-scribRNAi /+;* (O) *C885a-Gal4/UAS-Tak1^DN^; UAS-scribRNAi/+; (P) C885a-Gal4/UAS-Yki^S168A^; UAS-scribRNAi/+;* (Q) *C885a-Gal4/UAS-Ras^V12^; UAS-scribRNAi/+;* (R) *C885a-Gal4/UAS-NICD; UAS-scribRNAi/+.* Scale bar: 10μm. (S) Barplot of volumes for 4-day AEL control (n = 33, 1.0±0.3×10^6^μm^3^), *scribRNAi* (n = 30, 5±1×10^5^μm^3^)*, scribRNAi Tak1^DN^* (n = 35, 1.1±0.2×10^6^μm^3^)*, scribRNAi Yki^S168A^* (n = 26, 7±2 x10^5^μm^3^)*, scribRNAi Ras^V12^* (n = 30, 4±2×10^5^μm^3^), and *scribRNAi NICD* (n = 18, 2.1±0.4×10^5^μm^3^) imaginal discs. (T) Quantification of volumes for control, *scribRNAi, scribRNAi Tak1^DN^, scribRNAi Yki^S168A^, scribRNAi Ras^V12^,* and *scribRNAi NICD* imaginal discs over time. Control, **4d** n = 33, 1.0±0.3×10^6^μm^3^; **5d** n = 26, 2.1±0.6×10^6^μm^3^. *scribRNAi*, **4d** n = 30, 5±1×10^5^μm^3^; **5d** n = 27, 1.4±0.5×10^6^μm^3^; **6d** n = 34, 3±1×10^6^μm^3^; **7d** n = 34, 6±2×10^6^μm^3^; **8d** n = 32, 8±3×10^6^μm^3^. *scribRNAi Tak1^DN^*, **4d** n = 35, 1.1±0.2×10^6^μm^3^; **5d** n = 39, 3±1×10^6^μm^3^; **6d** n = 32, 6±3×10^6^μm^3^. *scribRNAi Yki^S168A^*, **4d** n = 26, 7±2×10^5^μm^3^; **5d** n = 30, 1.9±0.8×10^6^μm^3^; **6d** n = 32, 5±1×10^6^μm^3^; **7d** n = 28, 9±3×10^6^μm^3^; **8d** n = 34, 1.1±0.4×10^7^μm^3^. *scribRNAi Ras^V12^*, **4d** n = 30, 4±2×10^5^μm^3^; **5d** n = 35, 1.9±0.4×10^6^μm^3^; **6d** n = 30, 7±2×10^6^μm^3^; **7d** n = 28, 1.1±0.4×10^7^μm^3^; **8d** n = 28, 1.6±0.6×10^7^μm^3^. *scribRNAi NICD*, **4d** n = 18, 2.1±0.4×10^5^μm^3^; **5d** n = 20, 3±2×10^5^μm^3^; **6d** n = 21, 5±3×10^5^μm^3^; **7d** n = 18, 2.5±0.7×10^5^μm^3^; **8d** n = 25, 2.2±0.4×10^5^μm^3^.

Interestingly, the time-course analysis showed that several signaling pathways previously implicated in modulating the *scrib* mutant cell growth show quantitative changes in the *scrib* mutant tumors over time. For example, the *unpaired* (*upd*) genes, the JAK/STAT pathway ligands, are significantly regulated in the 8-day *scrib* mutant tumors in comparison with the wild type control, with fold changes consistent with those reported in a previous study (Figure S4A)(Bunker et al., 2015). Time-course analysis of the *upd* family gene transcription reveals a peak expression in the 5-day AEL *scrib* mutant tumors that decreases over time (Figure S4A). Meanwhile, the expression of *mirror*, which is repressed by JAK/STAT transcriptional activity (Zeidler et al., 1999), shows lowest expression level in 5-day AEL *scrib* mutant tumors that recovers over time (Figure S4A). Notch, EGFR, JNK and Hippo signaling activities have been previously shown to modulate the growth outcomes of the *scrib* mutant mosaic clones (Brumby and Richardson, 2003; Chen et al., 2012; Igaki et al., 2006; Pagliarini and Xu, 2003). Similar to the JAK/STAT pathway, a time-course analysis of well-established transcriptional targets of Notch, EGFR and JNK signaling pathways reveals quantitative changes during the *scrib* mutant tumor progression (Figure 3C-D, Figure S4). Therefore, it is likely that quantitative changes in these signaling activities over time determine the differential growth rates and cell cycle profiles we observed in early and late *scrib* mutant tumors.

In particular, the abnormally high JNK signaling activity or low Notch or low EGFR signaling activity could be the underlying reason for the slow growth phenotype we observed in early *scrib* mutant tumors. We found that the growth arrest phenotype in 4-day and 5-day AEL posterior *scrib* tumors were rescued through overexpression of a dominant-negative form of Basket (Bsk^DN^) (Igaki et al., 2006)(Figure 3E-L) or a dominant-negative form of Tak1 (Figure S5), which block JNK signaling activity. Overexpression of Ras^V12^ and Notch intracellular domain (NICD) in combination with *scrib* RNAi through *engrailed-*Gal4 caused lethality during embryogenesis or severe overall developmental delay even with a temperature-sensitive form of tubulin-Gal80. We therefore performed the growth analysis of *scrib* RNAi tumors in combination with overexpression of Ras^V12^ using a C885a-Gal4 which expresses early to induce tumorigenesis and does not cause overall developmental delay (Hrdlicka et al., 2002). We found that overexpression of Ras^V12^ or NICD or Yki^S168A^ (an active form of Yki) cannot rescue the slow growth phenotype in early *scrib* RNAi tumors (Figure 3M-S), even though overexpression of Ras^V12^ and Yki^S168A^ showed growth-promoting effects in later stages (Figure 3T). Note that overexpression of NICD with C885a-Gal4 still lead to larvae of overall small body size and therefore small tumor size (Figure 3T). Taken together, these data suggest that the slow growth phase in early *scrib* tumors is primarily caused by high JNK signaling activity.

### The *scrib* mutant tumors harbor heterogeneous cell populations of different proliferative states

The *scrib* mutant tumors display changes in growth rate and cell cycle profiles, as well as quantitative differences in activities of multiple signaling pathways over time. These data suggest that the *scrib* mutant tumors undergo a transition from a growth arrest state to a proliferative state. To gain further insights into this transition, we built a spatiotemporally-resolved transcriptomic landscape from staged *scrib* mutant tumors.

We profiled a minimum of 3000 cells per stage from 4-day, 5-day and 8-day AEL *scrib* mutant tumors. We then pooled the *scrib* mutant cells from different stages together for clustering and examined the distribution of four cell types defined by the expression levels of the JNK signaling activity reporter *Mmp1* and the EGFR signaling reporter *kek1*. *Mmp1^high^* cell number decreases and *kek1^high^* cell number increases over time, consistent with the bulk RNA-seq data and validating the scRNA data quality (Figure 4A-B). Moreover, *Mmp1* and *kek1* show significant biased distribution in single cells from different stages (Figure S6), suggesting that the *scrib* mutant tumors might harbor heterogeneous populations of cells at different proliferative states.

**Figure 4.**
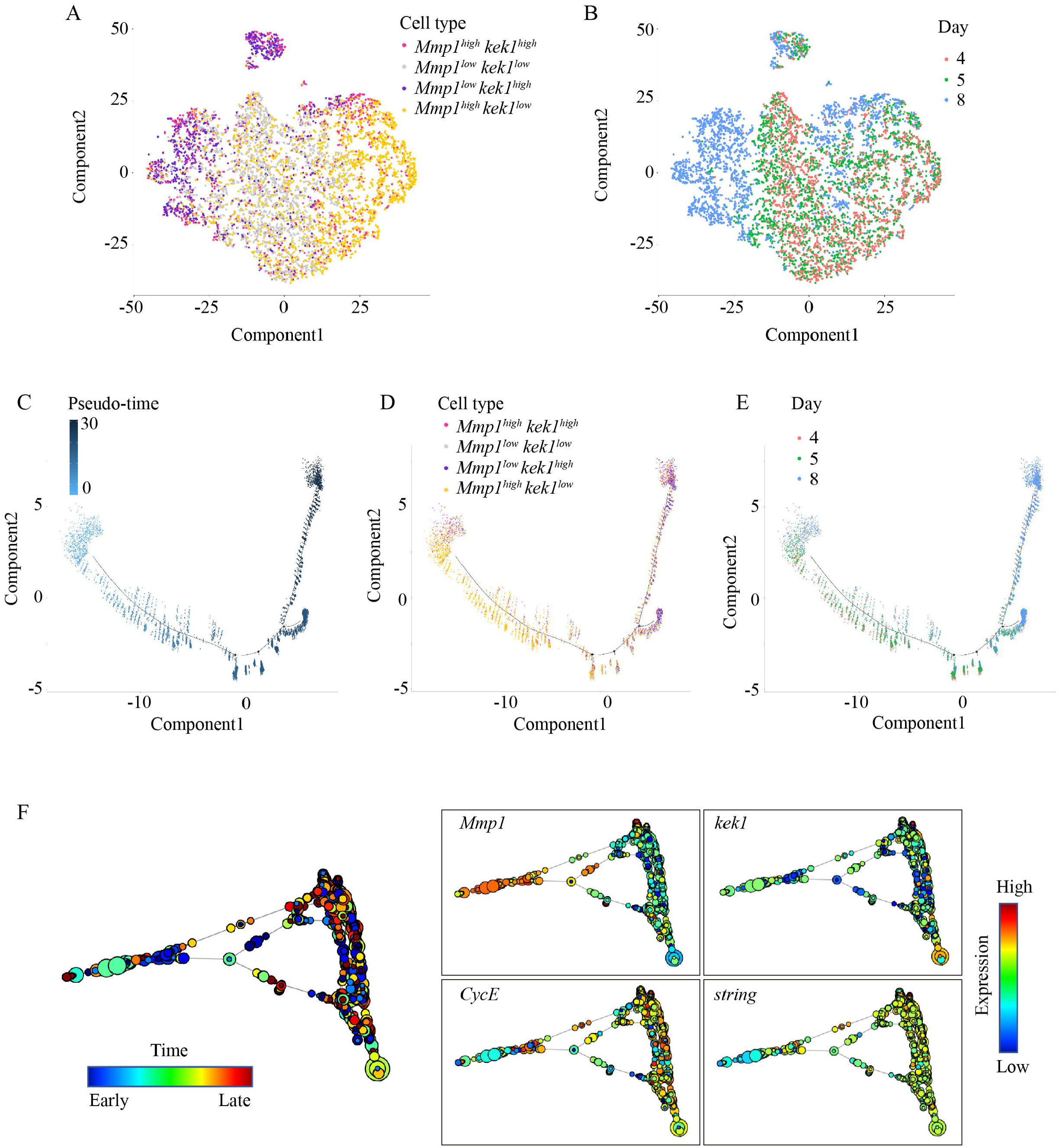
Visualization of single cells from the *scrib* mutant tumors in the reduced dimensional space. (A-B) Visualization of single cells ordered in the t-SNE reduced dimensional space colored by cell types defined by *Mmp1* and *kek1* expression level (A) (normalized expression level=1 as threshold), and tumor age (B). Single cells pooled from 4-day, 5-day and 8-day AEL *scrib* mutant tumors. 3000 single cells per time point are randomly sampled. (C-E) Visualization of single cells ordered along a pseudotime trajectory in the reduced dimensional space colored by pseudotime (C), cell types defined by *Mmp1* and *kek1* expression level (D) (normalized expression level=1 as threshold), and tumor age (E). 3000 single cells per time point are randomly sampled. (F) Topological representation of single cells ordered using scTDA. 500 cells per time point are randomly sampled. The nodes (circles) are clusters of single cells with similar global expression profiles, and the node size corresponds to the number of cells in that cluster. Edges (lines) connect clusters that have at least one cell in common. The node color in the left panel indicates the time line of the *scrib* mutant tumor progression. A node composed of a mixture of cells from early and late stages has an intermediate color. The node color in the right panel indicates the expression level of marker genes labeled in each panel.

To further explore the distribution of potential cell states, we first used *Mmp1* and *kek1* as marker genes to perform semi-supervised cell ordering in pseudotime with Monocle (Trapnell et al., 2014) (Figure 4C-E). The expression of JNK and EGFR signaling reporter genes show progressive changes along the pseudotime trajectory (Figure S7). Interestingly, the Hippo signaling activity reporters *Diap1*, *expanded(ex)* and *Cyclin E(CycE)* also show increase along the pseudotime trajectory (Figure S7), while these genes do not show obvious changes over time (Figure S6). Similarly, genes that promote cell cycle progression, such as *Cyclin B*, *Cdc25*/*string,* and *Cdc20/fizzy,* also show increase along the pseudotime trajectory (Figure S7). Taken together, the pseudotime trajectory likely reflects an arrest-to-proliferation state transition trajectory for the *scrib* mutant cells. We found that single cells from tumors of different ages are scattered along the pseudotime trajectory. Notably, the 4-day and 5-day AEL *scrib* mutant cells are more enriched towards the arrested state of the pseudotime trajectory and the 8-day AEL *scrib* mutant cells are enriched towards the opposite end along the trajectory (Figure 4E).

The above analysis is based on semi-supervised learning and the method assumes a treelike structure to distinguish cell states (Trapnell et al., 2014). To gain an unbiased view of the *scrib* mutant cell states, we further adopted the scTDA, a nonlinear, model-independent, and unsupervised topological data analysis (TDA) method for analyzing and visualizing single-cell data (Rizvi et al., 2017). We randomly selected the same number of cells from each stage and pooled these cells from all stages for analysis. Interestingly, different random experiments robustly capture well-clustered cells that likely represent the arrested state based on inspection of marker genes such as *Mmp1*, *kek1*, *CycE* and *string* (Figure 4F and Figure S8). Moreover, cells from different ages are scattered on the topological representation, again indicating that the *scrib* mutant tumors from all profiled stages harbor heterogeneous cell populations. Notably, the cells at the arrest state are more likely to be from the early stage tumors (Figure 4F and Figure S8). Interestingly, while the arrest state is well-clustered and easily detectable, other cell states form complex structures in the topological representations, indicating heterogeneous cell states presented in the *scrib* mutant tumors are unlikely to be along a linear transition from the least proliferative to the most proliferative state.

## Discussion

Here we demonstrated that the *scrib* mutant tumors undergo a transition from a growth-arrest state to a proliferation state over time and constructed a spatiotemporally resolved evolution landscape for the *scrib* mutant tumors. Our data suggest that the *scrib* mutant tumors harbor heterogeneous cell populations, providing a foundation for potential selection and transition into a proliferation state as a population over time.

We do not yet know how cells of different proliferative states arise in undifferentiated *scrib* mutant tumors. Notably, in flies a class of spindle assembly checkpoint mutations in combination with apoptosis blockage can induce neoplastic tumor growth (Morais da Silva et al., 2013). It is therefore interesting to speculate if the two types of neoplastic tumor growth might partially share a common basis of generating heterogeneous cells through genome instability. It is also possible that cells of different states arise from stochasticity and the differences are amplified through complex feedback loops within the cell-signaling network. It will be interesting to explore whether and how cell states defined by combinatorial signaling activities are passed to daughter cells through epigenetic markers. It will be also interesting to explore whether and how cells can transit among different proliferative states.

The clonal *scrib* mutant cells are eliminated through cell competition when they are surrounded by wild-type neighbors (Brumby and Richardson, 2003; Vaughen and Igaki, 2016; Yamamoto et al., 2017). It is noteworthy that the clonal *scrib* mutant cells are likely to be in a different state from *scrib* mutant cells in early homozygous tumors. Studies have shown overexpression of Ras^V12^, NICD and p35 can effectively block the clonal *scrib* mutant cells from apoptosis induced by cell competition (Brumby and Richardson, 2003; Pagliarini and Xu, 2003). In our study, we found that overexpression of Ras^V12^, NICD and p35 have little effects in relieving the early *scrib* mutant tumors from growth arrest. It will be interesting to profile the clonal *scrib* mutant cells in the future and compare how the clonal *scrib* mutant cells change cell state in response to cell competition signals.

## Experimental Procedures

### Fly stocks

The fly strains used in this study were: *scrib^1^* FRT82B/TM6B (Bilder et al., 2000), UAS-*scrib* RNAi on the 2^nd^ chromosome (Bloomington/BL38199), UAS-*scrib* RNAi on 3^rd^ chromosome (BL35748), UAS-*dlg* RNAi on the 3^rd^ chromosome (BL35772), y[1] v[1]; P{y[+t7.7]=CaryP}attP2 (BL36303, the 3rd chromosome TRiP line background strain), y[1] v[1]; P{y[+t7.7]=CaryP}attP40 (BL36304, the 2^nd^ chromosome TRiP line background strain), engrailed-Gal4 UAS-GFP (BL25752), engrailed-Gal4 (BL30564), UAS-p35 (Hay et al., 1994)(BL6298), Fly-FUCCI (BL55098), worGal4, UAS-cherry::Jupiter, Sqh::GFP (Cabernard and Doe, 2013), c885a-Gal4 (Hrdlicka et al., 2002) (BL6990), UAS-Ras^V12^ (Karim and Rubin, 1998), UAS-Yki^S168A^ (Oh and Irvine, 2009)(BL28818), UAS-NICD (Rebay et al., 1993), UAS-Bsk^DN^(Igaki et al., 2006)(a kind gift from Jose C Pastor-Pareja), UAS-Tak1^DN^ (BL58811).

### Immunohistochemistry

Around 50 embryos collected within 3 hours were put in an individual vial of fly food to avoid crowding and the larvae were raised at 25-degree incubator for appropriate lengths of time before dissection. Imaginal discs were fixed and stained according to standard protocols. The primary antibodies used were mouse anti-phospho-Histone3 (1:1000, Cell Signaling), rabbit anti-Dcp-1(1:50, Cell Signaling), rabbit anti-phospho-*drosophila* S6 (Romero-Pozuelo et al., 2017), goat anti-GFP (1:1000, Abcam) and rabbit anti-DsRed (1:500, Takara). The secondary antibodies conjugated with various Alexa Fluor dyes (ThermoFisher) were used at 1:500. Phalloidin conjugated with Alexa Fluor dyes (1:1000, ThermoFisher) and Hoechst (1:10000, ThermoFisher) were used to stain F-actin and DNA, respectively. For EdU incorporation assay, we labeled the dissected imaginal discs for 30 min before fixation using the Click-iT^TM^ Plus EdU Alexa Fluor^TM^594 Imaing Kit (ThermoFisher). All images were acquired on a Leica TCS SP8 confocal microscope.

### Western blotting

About 30 larvae were dissected in PBS. Cell lysates were homogenized in 1X RIPA (Millipore) with protease inhibitors (Roche). The primary antibodies used were mouse anti-MMP1 (1:100, DSHB) and mouse anti-alpha-tubulin (1:5000, DSHB).

### Image processing and data analysis

Images were taking as z-stacks with a step size of 1 μm. Tissue volume was measured with Measure Stack plugin in Fiji. PH3+ cell number was calculated with Cell Counter plugin in Fiji. EdU intensity were measured in unit areas from the posterior and anterior region respectively.

### Fluorescence-activated cell-sorting (FACS) analysis

Wing discs were dissected from staged larvae and dissociated for FACS analysis according to standard protocol (Neufeld et al., 1998). Cells were sorted with BDFACSAria IIIu and data were analyzed with FlowJo.

### *Drosophila* neuroblast live imaging

Female virgins of hsFLP; worGal4,Sqh::GFP,UASCherry::jupiter; FRT82B/TM6B were crossed with males of *scrib*FRT82B/TM6B. Progeny were heat shocked at 38 degree (in the water bath) for 1 hour and subsequently raised at 25 degree until imaging. Five-day AEL larvae were then dissected and imaged according to standard protocol (Cabernard and Doe, 2013). For wild type control, larvae expressing wt; worGal4,Sqh::GFP,UASCherry::jupiter; Dr/TM6B were dissected and imaged with the same laser setting as that of the *scrib* mutant neuroblasts.

### Bulk RNA-seq and data analysis

Total RNA was extracted from control and *scrib^1^* wing imaginal discs with RNeasy Mini Kit (Qiagen). Construction of cDNA libraries and 150bp paired-end sequencing on Illumina HiSeq platform were performed by Novogene. Cleaned raw reads were mapped to the reference genome using STAR and counts are generated by featuresCounts available in Subread package. PCA analysis was performed in DESeq2. Count normalization was performed using edgeR before hierarchical clustering (hclust function in R).

### 10x Genomics single cell RNA-seq and data analysis

Staged *scrib^1^* wing imaginal discs were dissected and transferred to DPBS (ThermoFisher). The wing imaginal discs were dissociated in 0.25% Trypsin-EDTA solution at 37 °C for 10 min. Cells were then washed in DPBS and passed through 35μm filter before library preparation. Construction of 10x single cell libraries and sequencing on Illumina Hiseq platform were performed by Novogene. Raw data mapping and primary analysis was performed in the Cell Ranger pipeline. Secondary analysis for marker gene expression pattern was performed with Cell Ranger R kit and Seurat. Cell type clustering and construction of single-cell pseudotime trajectory using semi-supervised DDRTree method was performed with Monocle (Trapnell et al., 2014). Unstructured and unsupervised topological data analysis (TDA) method was performed for ordering cells (Rizvi et al., 2017). To balance the cell number among stages, we randomly selected 500 cells per stage for each experiment and repeated the experiment for three times. Figures from different repeats were shown in Supplementary Figure 8.

## Author contributions

Y.Y., T.T.J. and L.Z. designed the experiments. T.T.J., L.Z. and Y.W. performed all the experiments except Figure S3. Y.Y., M.X.D. and T.T.J. analyzed all the data and prepared all the figures except Figure S3. S.S.H. and J.G.W provided RNA sequencing data analysis tools, performed TDA, prepared Figure 4F, Figure S6 and Figure S8. P.T., A.A.S, V.S and C.C. performed experiments for Figure S3 and prepared Figure S3. Y.Y. wrote the manuscript with input from J.G.W. and C.C.

## Acknowledgements

We thank Dr. Chris Doe and Dr. Jose C Pastor-Pareja, Bloomington Stock Center and Developmental Biology Hybridoma Bank for providing fly stocks and reagents. We thank Dr. Zilong Wen and Dr. Mingjie Zhang for sharing their confocal microscope. We thank Dr. Chris Doe and Dr. Trudi Schupbach for helpful comments on the manuscript. We thank Dr. Tatsushi Igaki for helpful discussion on the project. This work was supported by grants to Yan Yan from the Research Grants Council of the Hong Kong Special Administrative Region (grants GRF16103314, 16103815, 16150016, AoE/M-09/12) and to Jiguang Wang from from a NSFC/RGC Grant No. N_HKUST601/17 and a CRF Grant No. C6002-17G.

